# Social play behavior shapes the development of prefrontal inhibition in a region-specific manner

**DOI:** 10.1101/2023.02.13.528312

**Authors:** Ate Bijlsma, Louk J.M.J. Vanderschuren, Corette J. Wierenga

**Affiliations:** Department of Population Health Sciences, Division of Behavioural Neuroscience, Faculty of Veterinary Medicine, Utrecht University, Yalelaan 1, 3584 CL Utrecht, the Netherlands; Department of Biology, Faculty of Science, Utrecht University, Padualaan 8, 3584 CH Utrecht, the Netherlands; Donders Institute and Faculty of Science, Radboud University, Heyendaalseweg 135, 6525 AJ Nijmegen, the Netherlands

**Author notes:** **corresponding authors**, Prof. Dr. Corette J. Wierenga, Prof. Dr. Louk J.M.J. Vanderschuren. **Author Contribution:** The study was conceived by AB, LV and CW. Experiments were designed by AB, LV and CW. Experiments and analysis were performed by AB. Manuscript was written by AB, LV and CW.

**Keywords:** Brain development, Experience-dependent plasticity, Inhibitory signaling, Prefrontal cortex, Social play behavior

## Abstract

Experience-dependent organization of neuronal connectivity is critical for brain development. We recently demonstrated the importance of social play behavior for the developmental fine-tuning of inhibitory synapses in the medial prefrontal cortex (mPFC) in rats. When these effects of play experience exactly occur and if this happens uniformly throughout the prefrontal cortex is currently unclear. Here we report important temporal and regional heterogeneity in the impact of social play on the development of excitatory and inhibitory neurotransmission in the mPFC and the orbitofrontal cortex (OFC). We recorded in layer 5 pyramidal neurons from juvenile (postnatal day (P)21), adolescent (P42) and adult (P85) rats after social play deprivation (SPD; between P21-P42). The development of these PFC subregions followed different trajectories. On P21, inhibitory and excitatory synaptic input was multiple times higher in the OFC than in the mPFC. SPD did not affect excitatory currents, but reduced inhibitory transmission in both mPFC and OFC. Intriguingly, the reduction occurred in the mPFC during SPD, while the reduction in the OFC only became manifested after SPD. These data reveal a complex interaction between social play experience and the specific developmental trajectories of prefrontal subregions.

## Introduction

The developing brain requires proper external input to fine-tune activity and connectivity in neural circuits for optimal functionality throughout life. Experience-dependent plasticity is well described in the sensory cortex, but it is also essential for the development of higher-order brain regions, including the prefrontal cortex (PFC) (Larsen and Luna, 2018; Bicks et al., 2020). The PFC is undergoing intensive functional remodeling during the juvenile and adolescent phases of life, roughly between postnatal day (P) 21 and 85 in rodents (Kolb et al., 2012; Thomases et al., 2013; Caballero and Tseng, 2016; Caballero et al., 2016; Larsen and Luna, 2018). Cytoarchitectonic characteristics of the PFC do not stabilize until around P30. Around this time, white matter volume increases because of myelination, and grey matter volume in the PFC starts to decrease due to synaptic pruning and apoptosis (Markham et al., 2007). In addition, at the onset of adolescence (around P30 – P42), the PFC starts to receive long-range afferents from sensory and subcortical brain regions including the amygdala, ventral hippocampus and mediodorsal thalamus (Hoover and Vertes, 2007, 2011; Murphy and Deutch, 2018; Yang et al., 2021). During adolescence (P35-P60), local interneurons are undergoing important remodeling (Caballero et al., 2014a; Cass et al., 2014; Caballero and Tseng, 2016), which is critical for the maturation of the PFC network (Tseng et al., 2008). Proper functioning of the prefrontal network in adulthood is important for working memory and cognitive flexibility (Murray et al., 2015) and the vulnerability to fear (Courtin et al., 2014), stress (Sriparna Ghosal, Brendan Hare, 2017) and psychosis (Tanaka, 2008).

During the juvenile and adolescent phases of life, when PFC development is in progress, most mammalians species, including rats and humans, display an abundance of a pleasurable and energetic form of social interaction, known as social play behavior (Panksepp et al., 1984; Vanderschuren et al., 1997; Pellis and Pellis, 2009; Manduca et al., 2014). One important characteristic of social play is that it allows animals to experiment with their own behavior and their interactions with others. This experimentation during social play is thought to facilitate the development of a rich behavioral repertoire, that allows an individual to quickly adapt in a changeable world. In this way, social play may subserve the development of PFC-dependent skills such as flexibility, creativity, and decision-making (Špinka et al., 2001; Pellis and Pellis, 2009; Vanderschuren and Trezza, 2014). Indeed, during play the PFC is engaged (Van Kerkhof et al., 2014) and required (Bell et al., 2009; van Kerkhof et al., 2013). Moreover, limiting the time young animals can play has been shown to lead to impaired social interactions (Hol et al., 1999; Van Den Berg et al., 1999) and long-lasting changes in PFC function and circuitry in adulthood (Bell et al., 2010; Baarendse et al., 2013; Vanderschuren and Trezza, 2014).

The PFC comprises multiple subregions that display functional specialization and overlap (Miller and Cohen, 2001; Dalley et al., 2004; Izquierdo et al., 2017; Verharen et al., 2020). Of these, both the medial prefrontal cortex (mPFC) and the orbitofrontal cortex (OFC) are required for social play (Schneider and Koch, 2005; Pellis et al., 2006; Bell et al., 2009; Van Kerkhof et al., 2013), and social play facilitates the maturation of these regions (Pellis et al., 2010; Baarendse et al., 2013; Himmler et al., 2018). Both subregions have been implicated in higher cognitive, so-called executive functions, whereby the OFC is thought to have the upper hand in emotionally colored cognition, such as reward-based decision making (Schoenbaum et al., 1998, 2009; Rolls, 2000; O’Doherty et al., 2003), and the mPFC subserves functions in working memory and planning (Bechara and Damasio, 2005; Posner et al., 2007; Euston et al., 2012). However, there is also a substantial degree of functional overlap between PFC regions (Sul et al., 2010; Lodge, 2011; Hardung et al., 2017). During the production of social behaviors the mPFC and OFC are functionally linked to each other and the two regions are reciprocally connected (Singer et al., 2009; Hoover and Vertes, 2011).

We recently showed that deprivation of social play affects inhibitory, but not excitatory connections in the adult mPFC, emphasizing the importance of social play for PFC circuit development (Bijlsma et al., 2022). How social play contributes to the development of OFC connections is currently unknown. Additionally, how the excitatory and inhibitory inputs of the two subregions develop and how the deprivation of social play experiences affects their developmental trajectories has not been addressed. Here we report the distinct development of synaptic inputs onto layer 5 pyramidal neurons in the mPFC and OFC and describe how social play deprivation (SPD) differentially affects the developmental trajectories of these two PFC regions.

## Materials and Methods

### Animals and housing conditions

All experimental procedures were approved by the Animal Ethics Committee of Utrecht University and the Dutch Central Animal Testing Committee and were conducted in accordance with Dutch (Wet op de Dierproeven, 1996; Herziene Wet op de Dierproeven, 2014) and European legislation (Guideline 86/609/EEC; Directive 2010/63/EU). Male Lister Hooded rats were obtained from Charles River (Germany) on postnatal day (P) 14 in litters with nursing mothers. All rats were subject to a normal 12:12h light-dark cycle with ad libitum access to water and food. Rats used in the P21 measurements were directly taken from the litter at P21. Rats used in the P42 and P85 groups were weaned on P21 and were either allocated to the control (CTL) group or the social play deprivation (SPD) group. CTL rats were housed in pairs for the remainder of the experiment. SPD rats were pair-housed but during P21 to P42 a transparent Plexiglas divider containing small holes was placed in the middle of their home cage creating two separate, identical compartments. SPD rats were able to see, smell and hear one another but they were unable to physically interact. On P42, the Plexiglas divider was removed and SPD rats were housed in pairs for the remainder of the experiment. Rats were weighed and handled at least once a week until they were used for neurophysiological experiments. Experiments were performed on P21, P42 and P85 with a spread of 2 days as it was not always possible to perform measurements on the exact postnatal day.

### Electrophysiological analysis

*Slice preparation:* Rats were anaesthetized with Isoflurane and then transcardially perfused with ice-cold modified artificial cerebrospinal fluid (ACSF) containing (in mM): 92 Choline chloride, 2.5 KCl, 1.2 NaH_2_PO_4_, 30 NaHCO_3_, 20 HEPES, 25 glucose, 5 Na-ascorbate, 3 Na-pyruvate, 0.5 CaCl_2_, and 10 MgSO_4_, bubbled with 95% O_2_ and 5% CO_2_ (pH 7.3–7.4). The brain was quickly removed after decapitation and coronal slices (300 μm) of the medial PFC (consisting of the prelimbic and infralimbic cortex) and OFC (consisting of the ventral and lateral orbital cortex) were prepared using a vibratome (Leica VT1000S, Leica Microsystems) in ice-cold modified ACSF. Slices were initially incubated in the carbonated modified ACSF for 5 min at 35 °C and then transferred into a holding chamber containing standard ACSF containing (in mM): 126 NaCl, 3 KCl, 1.3 MgCl_2_, 2 CaCl_2_.2H2O, 20 glucose, 1.25 NaH_2_PO_4_ and 26 NaHCO_3_ bubbled with 95% O_2_ and 5% CO_2_ (pH 7.3) at room temperature for at least 30 minutes. They were subsequently transferred to the recording chamber, perfused with standard ACSF that is continuously bubbled with 95% O_2_ and 5% CO_2_ at 28–32 °C. *Whole-cell recordings and analysis:* Whole-cell patch-clamp recordings were performed from layer 5 pyramidal neurons in the medial PFC and OFC. Neurons were visualized with an Olympus BX51W1 microscope using infrared video microscopy and differential interference contrast (DIC) optics. Patch electrodes were pulled from borosilicate glass capillaries and had a resistance of 4-6 MΩ when filled with intracellular solutions. Excitatory postsynaptic currents (EPSCs) were recorded with an internal solution containing (in mM): 140 K-gluconate, 4 KCl, 10 HEPES, 0.5 EGTA, 4 MgATP, 0.4 NaGTP, 4 Na_2_-phosphocreatine (pH 7.3 with KOH). Spontaneous inhibitory postsynaptic currents (sIPSCs) were recorded in the presence of 6,7-dinitroquinoxaline-2,3-dione (DNQX) (20 μM) and D,L-2-amino-5-phosphopentanoic acid (D,L-AP5) (50 μM), with an internal solution containing (in mM): 70 K-gluconate, 70 KCl, 10 HEPES, 0.5 EGTA, 4 MgATP, 0.4 NaGTP, 4 Na_2_-phosphocreatine (pH 7.3 with KOH). Action-potential independent miniature IPSCs (mIPSCs) were recorded under the same conditions as sIPSCs, but in the presence of 1 μM tetrodotoxin (TTX) to block voltage-gated sodium channels. The membrane potential was held at −70 mV for voltage-clamp experiments. Signals were amplified, filtered at 2 kHz and digitized at 10 kHz using a MultiClamp 700B amplifier (Molecular Devices) and stored using pClamp 10 software. Series resistance was constantly monitored, and the cells were rejected from analysis if the resistance changed by >20% during the experiment or reached a value higher than 30 MΩ. No series resistance compensation was used. Resting membrane potential was measured in bridge mode (I=0) immediately after obtaining whole-cell access. The basic electrophysiological properties of the cells were determined from the voltage responses to a series of hyperpolarizing and depolarizing square current pulses. Passive and active membrane properties were analysed with Matlab (R2019b, MathWorks) using a custom script. Miniature and spontaneous synaptic currents (IPSCs and EPSCs) data were analysed with Mini Analysis (Synaptosoft). The detected currents were manually inspected to exclude false events.

### Data processing and statistical analyses

Statistical analyses and data processing were performed with GraphPad Prism (Software Inc.) and RStudio 1_2_5019 (R version 3.6.1, R Foundation for Statistical Computing). The variance between cells within slices was larger than the variance between slices, indicating that individual cells can be treated as independent measurements. Differences between time points (P21, P42 and P85) were tested with one-way ANOVA followed by a Tukey’s test when significant (denoted in figures by a color-coded asterisk in blue for CTL and red for SPD). Differences between groups were tested with two-way ANOVA followed by a Tukey’s test (black asterisks in figures). Percentage growth for the P21-P42 timeframe was calculated by normalizing the values of P42 (CTL and SPD) to the mean of P21. For the P42-P85 timeframe, the P85 values were normalized to the P42 mean of the same condition. All graphs represent the mean ± standard error of the mean (SEM) with individual data points shown in colored circles.

## Results

We performed whole cell patch clamp recordings in layer 5 (L5) pyramidal cells in the mPFC (Fig. 1A) in slices prepared from juvenile (P21), adolescent (P42) and adult (P85) control (CTL) male rats to assess the development of their synaptic input currents (Fig. 1B,C). We found that the frequency of inhibitory inputs onto L5 mPFC pyramidal neurons strongly increased between P21 and P85. A large, 3-fold, increase in sIPSC frequency occurred between P21 and P42 (Fig 1D,E). Between P42 and P85, a smaller ~60% increase in sIPSC frequency was observed, while large individual differences between L5 cells emerged (Fig 1D,E). Amplitudes of the sIPSCs remained stable across time points (Fig. 1F), while rise and decay kinetics were faster at P42 compared to P21, an effect that was less prominent at P85 (Fig. 1G,H). This suggests that the inhibitory synaptic inputs to L5 cells in the mPFC are undergoing intense development between P21 and P42, with a smaller rate of growth after P42 until adulthood. These findings are in agreement with previous studies showing an increase in inhibitory synaptic inputs (Cass et al., 2014; Kalemaki et al., 2020) and accelerating kinetics (Vicini et al., 2001; Hashimoto et al., 2010) during early development.

**Fig. 1.**
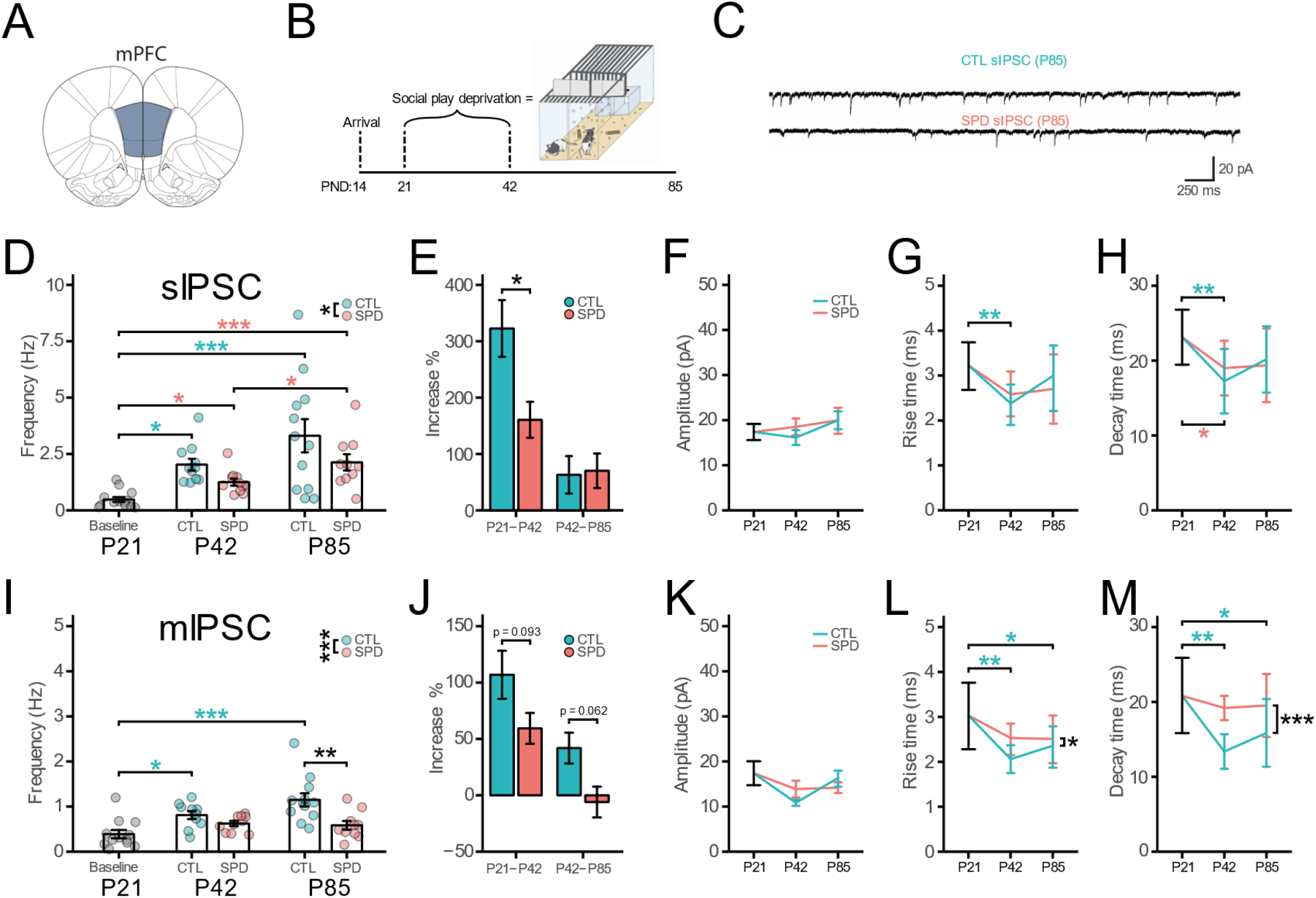
(**A**) Schematic diagram depicting the recording site in the mFPC. (**B**) Social play deprivation (SPD) paradigm. (**C**) Example traces of spontaneous inhibitory postsynaptic currents (sIPSCs) in L5 pyramidal cells in slices from P85 control (CTL) and SPD rats. (**D**) Frequency of sIPSCs in Baseline (P21), CTL and SPD slices (P42 & P85) (CTL 1W-ANOVA, Time: p<0.001; SPD 1W-ANOVA, Time: p<0.001; 2W-ANOVA, Condition: p=0.031, Time: p=0.023, Interaction: p=0.67). (**E**) Percentage increase of sIPSC frequency from P21 to P42 and P42 to P85 for both CTL and SPD slices (P21-P42 T-Test, p=0.018; P42-P85 T-Test, p=0.88). (**F-H**) Amplitude (**F**) (CTL 1W-ANOVA, Time: p=0.34; SPD 1W-ANOVA, Time: p=0.72; 2W-ANOVA, Condition: p=0.63, Time: p=0.22, Interaction: p=0.56), Rise time (**G**) (CTL 1W-ANOVA, Time: p=0.005; SPD 1W-ANOVA, Time: p=0.045; 2W-ANOVA, Condition: p=0.95. Time: p=0.078, Interaction: p=0.24) and Decay time (**H**) (CTL 1W-ANOVA, Time: p=0.004; SPD 1W-ANOVA, Time: p=0.020; 2W-ANOVA, Condition: p=0.77, Time: p=0.23, Interaction: p=0.36) of sIPSC events. (**I**) Frequency of miniature inhibitory postsynaptic currents (mIPSCs) in Baseline (P21), CTL and SPD slices (P42 & P85) (CTL 1W-ANOVA, Time: p<0.001; SPD 1W-ANOVA, Time: p<0.001; 2W-ANOVA, Condition: p<0.001, Time: p=0.15, Interaction: p=0.089) (**J**) Percentage increase of mIPSC frequency from P21 to P42 and P42 to P85 for both CTL and SPD slices (P21-P42 T-Test, p=0.093; P42-P85 T-Test, p=0.062). (**K-M**) Amplitude (**K**) (CTL 1W-ANOVA, Time: p=0.072; SPD 1W-ANOVA, Time: p=0.42; 2W-ANOVA, Condition: p=0.85, Time: p=0.058, Interaction: p=0.10), Rise time (**L**) (CTL 1W-ANOVA, Time: p=0.001; SPD 1W-ANOVA, Time: p=0.081; 2W-ANOVA, Condition: p=0.042, Time: p=0.32, Interaction: p=0.34) and Decay time (**M**) (CTL 1W-ANOVA, Time: p<0.001; SPD 1W-ANOVA, Time: p=0.61; 2W-ANOVA, Condition: p<0.001, Time: p=0.20, Interaction: p=0.34) of mIPSC events. (**D-H**) Data from 13 (P21), 11 (P42 CTL), 11 (P42 SPD), 12 (P85 CTL), 10 (P85 SPD) cells. (**I-M**) Data from 12 (P21), 10 (P42 CTL), 10 (P42 SPD), 12 (P85 CTL), 10 (P85 SPD) cells. Statistical range: * p≤0.05; ** p<0.01; *** p<0.001

We previously showed that SPD during P21-42 results in a reduction of inhibitory synapses onto L5 pyramidal somata in the mPFC of adult rats (Bijlsma et al., 2022). Here we assessed how SPD (Fig. 1B) affects the developmental trajectory of the synaptic circuitry in the mPFC. We observed that the large increase in sIPSCs found in CTL animals between P21 and P42 was reduced in SPD animals, and sIPSC frequency modestly increased between P42 to P85 (Fig. 1D). Interestingly, when the developmental increase was calculated relative to the sIPSC frequency at P42, we observed that the sIPSC reduction in L5 cells was entirely attributable to the SPD period between P21 and P42, while the increase from P42 to P85 was comparable in both conditions (Fig. 1E). SPD did not affect sIPSC amplitude (Fig. 1F) and the developmental acceleration of rise and decay time (Fig. 1G,H).

We also recorded miniature inhibitory postsynaptic currents (mIPSCs) in the presence of TTX which blocked all neuronal activity in the slices. In CTL slices, mIPSC frequency doubled between P21 and P42, followed by a smaller increase between P42 and P85 (Fig. 1I,J). Consistent with our observations for sIPSCs, mIPSC amplitudes did not change over this developmental period (Fig. 1K), while rise and decay kinetics became faster (Fig. 1L,M). The developmental increase in the frequency of mIPSCs was smaller compared to sIPSCs (compare fig. 1E and 1J), which suggests that the increase in inhibitory currents reflects the formation of new inhibitory synapses during this period as well as an increase in activity-dependent release. Consistent with our previous findings (Bijlsma et al., 2022), mIPSC frequency was reduced in the mPFC of SPD slices at P85, but the reduction was less pronounced at P42 (Fig. 1I). The developmental gain in mIPSC frequency between P21 and P42 was weaker in SPD rats compared to CTL rats and mIPSC frequency remained stable between P42 and P85, while mIPSC frequency in CTL rats still increased (Fig. 1J). The amplitude (Fig. 1K) of the mIPSCs were not affected by SPD, but the acceleration of rise and decay kinetics appeared less pronounced compared to CTL (Fig. 1L,M). Together, these results indicate that SPD interferes with the development of activity-dependent and -independent inhibitory currents in L5 cells of the mPFC and that the strongest effect is observed immediately after the deprivation period.

We previously showed that excitatory synaptic currents in L5 cells were unaffected by SPD in the adult mPFC. However, SPD may influence the developmental time course of excitatory synapse formation in the mPFC. We therefore measured excitatory synaptic inputs in mPFC slices from CTL and SPD rats at all three ages. In CTL slices, we observed a large increase in sEPSC frequency between P21 and P42, but sEPSC frequency remained stable after P42 (Fig. 2A,B). This is in line with the reported developmental increase of sEPSCs onto L5 fast-spiking interneurons in mPFC slices in a similar developmental period (Caballero et al., 2014a). The amplitudes of excitatory inputs remained stable over this period (Fig. 2C). The rise and decay times of sEPSCs were comparable between all time points (Fig. 2D,E). SPD did not affect any aspect of sEPSCs (Fig. 2A-E). These data indicate that similar to inhibitory synapses, excitatory synaptic inputs to L5 neurons in the mPFC undergo strong growth between P21 and P42. However, in stark contrast to inhibitory synapses, the development of excitatory synapses is not affected by SPD.

**Fig. 2.**
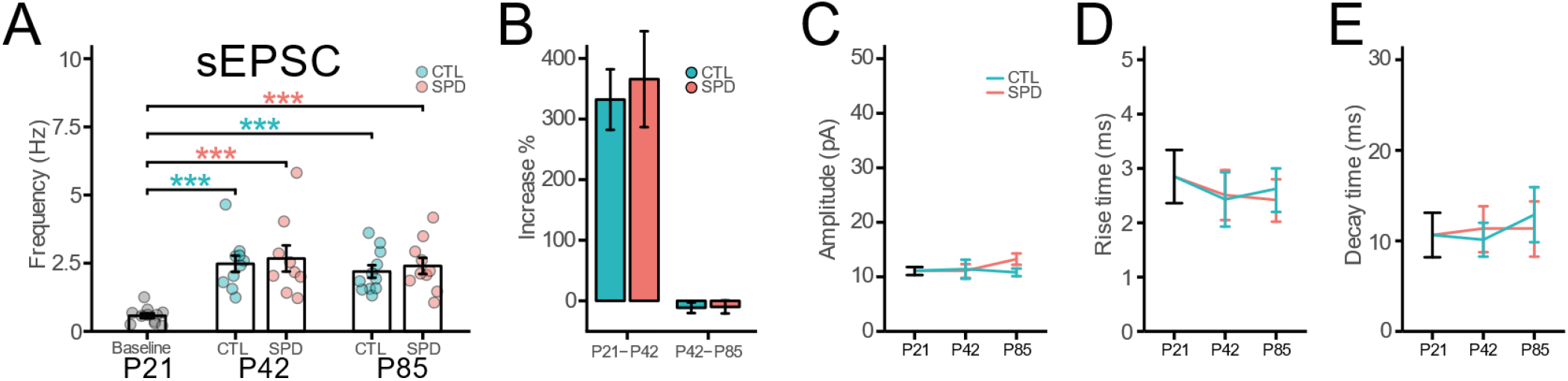
(**A**) Frequency of spontaneous excitatory postsynaptic currents (sEPSCs) in Baseline (P21), CTL and SPD slices (P42 & P85) (CTL 1W-ANOVA, Time: p<0.001; SPD 1W-ANOVA, Time: p<0.001; 2W-ANOVA, Condition: p=0.55, Time: p=0.40, Interaction: p=0.99). (**B**) Percentage increase of sEPSC frequency from P21 to P42 and P42 to P85 for both CTL and SPD mPFC slices (P21-P42 T-Test, p=0.73; P42-P85 T-Test, p=0.93). (**C-E**) Amplitude (**C**) (CTL 1W-ANOVA, Time: p=0.94; SPD 1W-ANOVA, Time: p=0.22; 2W-ANOVA, Condition: p=0.36, Time: p=0.55, Interaction: p=0.27), Rise time (**D**) (CTL 1W-ANOVA, Time: p=0.15; SPD 1W-ANOVA, Time: p=0.11; 2W-ANOVA, Condition: p=0.70, Time: p=0.76, Interaction: p=0.38) and Decay time (**E**) (CTL 1W-ANOVA, Time: p=0.052; SPD 1W-ANOVA, Time: p=0.84; 2W-ANOVA, Condition: p=0.76, Time: p=0.11, Interaction: p=0.14) of sEPSC events. (**A-E**) Data from 11 (P21), 10 (P42 CTL), 9 (P42 SPD), 11 (P85 CTL), 10 (P85 SPD) cells. Statistical range: * p≤0.05; ** p<0.01; *** p<0.001

We also assessed the intrinsic excitability of L5 pyramidal neurons in mPFC slices from CTL and SPD rats. We recorded action potentials (APs) during a series of increasing current injections. We observed that the intrinsic excitability of CTL cells slightly decreased from P21 to P42 and this was maintained in the P85 rats (Fig. 3A). AP threshold remained stable between P21 and P42 but was slightly lower at P85 (Fig. 3B). The membrane potential (Fig. 3C) and input resistance (Fig. 3D) were not different between time points. In slices from SPD rats, the developmental reduction in intrinsic excitability between P21 and P42 (Fig. 3E) was comparable to the CTL animals (comparing Fig. 3A and Fig. 3E, P42 CTL-SPD 2W-ANOVA, condition: p=0.097). This reduction was partly reversed, especially at lower current injections, in P85 rats. When comparing CTL and SPD cells at P85, no differences were found in AP number (comparing Fig. 3A and Fig. 3E, P85 CTL-SPD 2W-ANOVA, condition: p=0.79). The AP threshold (Fig. 3B), resting membrane potential (Fig. 3C) and input resistance (Fig. 3D) of the recorded neurons remained unaffected by SPD, in line with current literature (Baarendse et al., 2013; Bicks et al., 2020; Yamamuro et al., 2020). Our data indicate that AP firing in L5 pyramidal neurons is slightly reduced over development and that this is only mildly affected by SPD.

**Fig. 3.**
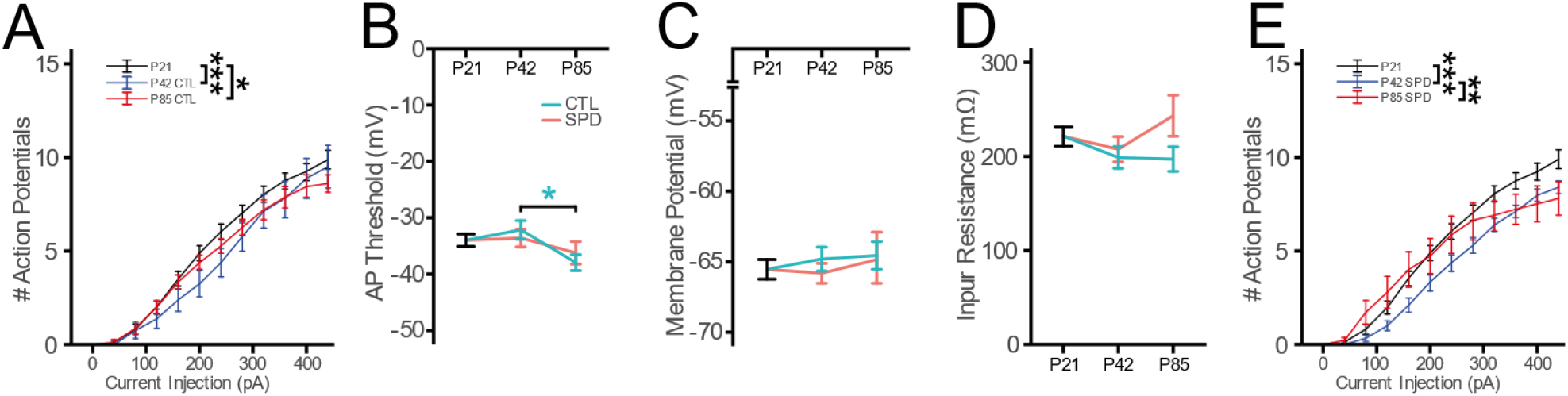
(**A**) Number of action potentials after current injections in Baseline (P21) and CTL (P42 & P85) mPFC slices (2W-ANOVA, AP: p<0.001, Current: p<0.001, Interaction: p= 0.93). (**B-D**) AP Threshold (**B**) (CTL 1W-ANOVA, Time: p=0.039; SPD 1W-ANOVA, Time: p=0.51; 2W-ANOVA, Condition: p=0.86, Time: p=0.020. Interaction: p=0.36), resting potential (**C**) (CTL 1W-ANOVA, Time: p=0.68; SPD 1W-ANOVA, Time: p=0.77; 2W-ANOVA, Condition: p=0.55, Time: p=0.53, Interaction: p=0.70) and input resistance (**D**) (CTL 1W-ANOVA, Time: p=0.26; SPD 1W-ANOVA, Time: p=0.31; 2W-ANOVA, Condition: p=0.13. Time: p=0.26, Interaction: p=0.24). (**E**) Number of action potentials after current injections in Baseline (P21) and SPD (P42 & P85) slices (2W-ANOVA, AP: p<0.001, Current: p<0.001, Interaction: p= 0.030). (**A-E**) Data from 33 (P21), 16 (P42 CTL), 21 (P42 SPD), 15 (P85 CTL), 14 (P85 SPD) cells. Statistical range: * p≤0.05; ** p<0.01; *** p<0.001

In contrast to the mPFC, developmental studies on circuitry development in the OFC are scarce or even absent. We therefore compared the development of the OFC and the mPFC and we assessed how SPD affects OFC development. Interestingly, the frequency of sIPSCs on OFC L5 pyramidal neurons at P21 (Fig. 4A,B) was 6-fold higher than in mPFC P21 slices in CTL rats. The sIPSC frequency remained stable between P21 and P42, and between P42 and P85 the frequency doubled (Fig. 4C,D). Similar to the mPFC, sIPSC amplitudes remained stable across time points (Fig. 4E) while the rise and decay kinetics became slightly faster in P42 and P85 rats compared to juvenile animals (Fig. 4F,G). This suggests that inhibitory synaptic inputs to L5 cells in the OFC only undergo growth after P42.

**Fig. 4.**
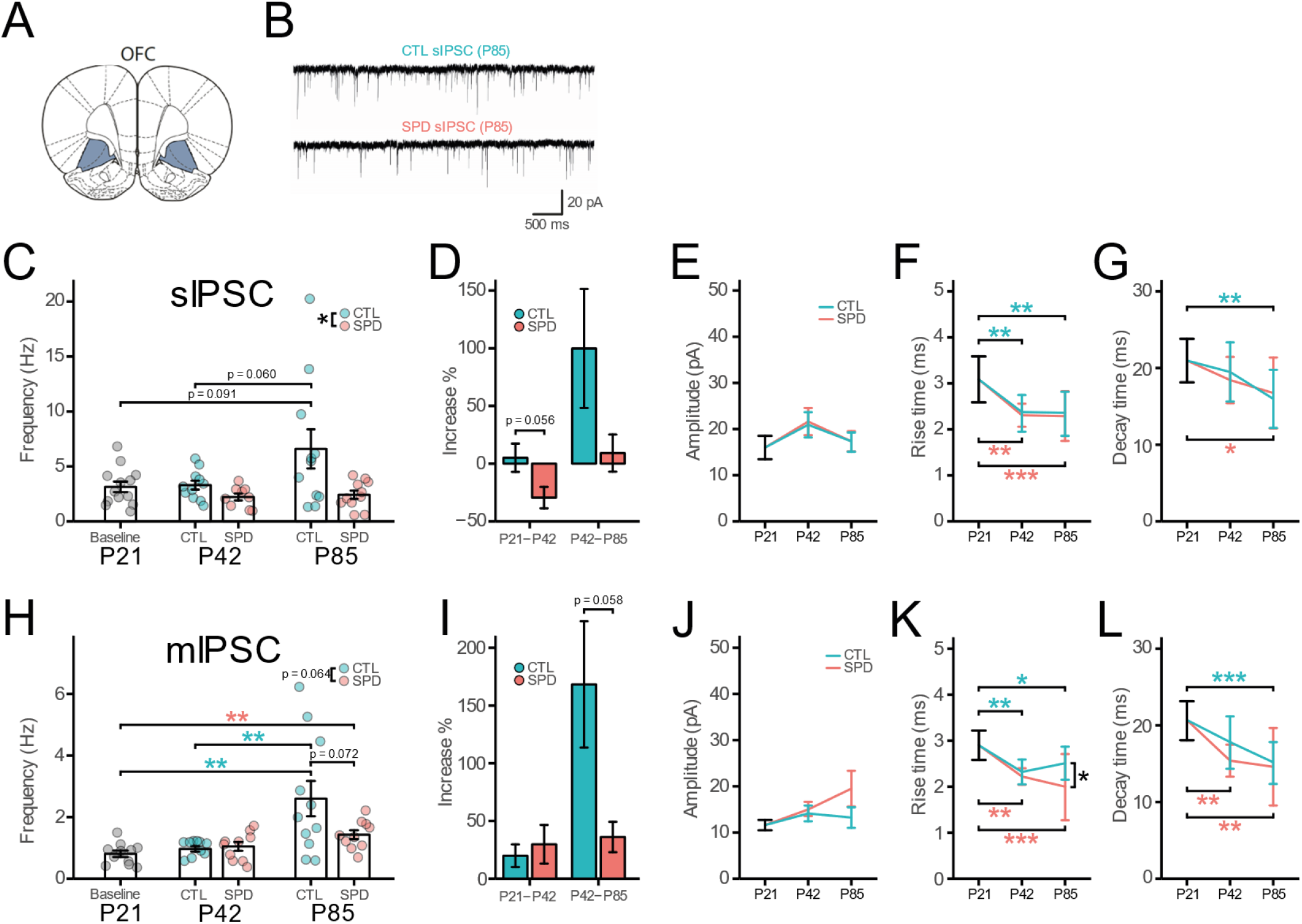
(**A**) Schematic diagram depicting the recording site in the OFC. (**B**) Example traces of spontaneous inhibitory postsynaptic currents (sIPSCs) in L5 pyramidal cells in slices from P85 control (CTL) and SPD rats. (**C**) Frequency of sIPSCs in Baseline (P21), CTL and SPD slices (P42 & P85) (CTL 1W-ANOVA, Time: p=0.045; SPD 1W-ANOVA, Time: p=0.26; 2W-ANOVA, Condition: p=0.012, Time: p=0.073, Interaction: p=0.13). (**D**) Percentage increase of sIPSC frequency from P21 to P42 and P42 to P85 for both CTL and SPD slices (P21-P42 T-Test, p=0.056; P42-P85 T-Test, p=0.12). (**E-G**) Amplitude (**E**) CTL 1W-ANOVA, Time: p=0.35; SPD 1W-ANOVA, Time: p=0.310; 2W-ANOVA, Condition: p=0.93, Time: p=0.12, Interaction: p=0.92), Rise time (**F**) (CTL 1W-ANOVA, Time: p<0.001; SPD 1W-ANOVA, Time: p<0.001; 2W-ANOVA, Condition: p=0.74, Time: p=0.95, Interaction: p=0.97) and Decay time (**G**) (CTL 1W-ANOVA, Time: p=0.007; SPD 1W-ANOVA, Time: p=0.032; 2W-ANOVA, Condition: p=0.85, Time: p=0.044, Interaction: p=0.47) of sIPSC events. (**H**) Frequency of miniature inhibitory postsynaptic currents (mIPSCs) in Baseline (P21), CTL and SPD slices (P42 & P85) (CTL 1W-ANOVA, Time: p=0.002; SPD 1W-ANOVA, Time: p=0.008; 2W-ANOVA, Condition: p=0.064, Time: p=0.006, Interaction: p=0.077). (**I**) Percentage increase of mIPSC frequency from P21 to P42 and P42 to P85 for both CTL and SPD slices (P21-P42 T-Test, p=0.76; P42-P85 T-Test, p=0.058). (**J-L**) Amplitude (**J**) (CTL 1W-ANOVA, Time: p=0.51; SPD 1W-ANOVA, Time: p=0.084; 2W-ANOVA, Condition: p=0.17, Time: p=0.53, Interaction: p=0.27), Rise time (**K**) (CTL 1W-ANOVA, Time: p=0.002; SPD 1W-ANOVA, Time: p<0.001; 2W-ANOVA, Condition: p=0.035, Time: p=0.89, Interaction: p=0.16) and Decay time (**L**) (CTL 1W-ANOVA, Time: p<0.001; SPD 1W-ANOVA, Time: p=0.001; 2W-ANOVA, Condition: p=0.27, Time: p=0.15, Interaction: p=0.43) of mIPSC events. (C-G) Data from 13 (P21), 11 (P42 CTL), 9 (P42 SPD), 11 (P85 CTL), 11 (P85 SPD) cells. (**E-L**) Data from 11 (P21), 9 (P42 CTL), 10 (P42 SPD), 11 (P85 CTL), 10 (P85 SPD) cells Statistical range: * p≤0.05; ** p<0.01; *** p<0.001

Similar to the mPFC, sIPSC frequency in the OFC in slices from SPD rats was reduced compared to CTL at P85. The sIPSC frequency in SPD slices showed a small reduction between P21 and P42, and the increase that was observed in CTL rats between P42 and P85 was completely absent in SPD rats (Fig. 4C,D). Similar to our observations in the mPFC, SPD did not affect sIPSC amplitude (Fig. 4E) or the acceleration in rise and decay kinetics (Fig. 4F, G). These results show that SPD reduces sIPSC frequency slightly during the deprivation period and impairs the growth of inhibitory synapses onto L5 neurons in the OFC afterwards.

The developmental time course of mIPSCs was comparable to that of sIPSCs, with no change in frequency between P21 and P42 and a ~2-fold increase between P42 and P85 (Fig. 4H,I). Event amplitudes remained unchanged across the different time points (Fig. 4J) with both rise and decay kinetics becoming faster after P21 (Fig. 4K,L), similar to the mPFC. We did not find any effect of SPD on the mIPSC frequency between P21 and P42 (Fig. 4H), while the increase between P42 and P85 was substantially reduced after SPD compared to CTL (Fig. 4I). SPD did not affect mIPSC amplitude (Fig. 4J), rise (Fig. 4K) and decay time (Fig. 4L). As the frequency of sIPSCs was substantially higher than of mIPSCs at P21, it is clear that there was already an activity-dependent component in the sIPSCs in the OFC at this early age, which was different from the mPFC. The effects of SPD are comparable between spontaneous and miniature inhibitory currents, suggesting that the SPD effect does not depend on activity but reflects a reduction in the number of inhibitory synapses after P42.

We also assessed the development of excitatory synapses in the OFC. Similar to the inhibitory synapses, sEPSC frequency was higher at P21 in the OFC compared to the mPFC. sEPSC frequency decreased between P21 and P42 and then remained stable until P85 (Fig. 5A,B). Amplitudes showed a small increase between P42 and P85 (Fig. 5C). sEPSC rise kinetics became faster with development (Fig. 5D), while the decay kinetics remained stable (Fig. 5E). SPD rats showed a similar decrease in sEPSC frequency between P21 and P42 and an increase in sEPSC amplitude between P42 and P85 compared to CTL rats. No differences were found in rise and decay kinetics after SPD (Fig. 5A-E). This indicates that similar to the mPFC, SPD did not affect the development of excitatory currents in the OFC.

**Fig. 5.**
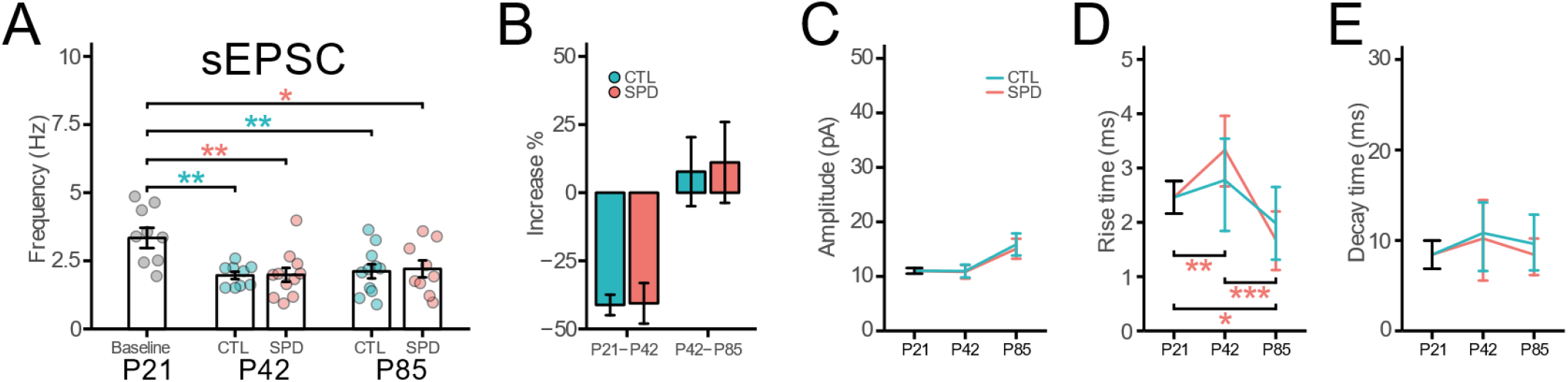
(**A**) Frequency of spontaneous excitatory postsynaptic currents (sEPSCs) in Baseline (P21), CTL and SPD OFC slices (P42 & P85) (CTL 1W-ANOVA, Time: p=0.002; SPD 1W-ANOVA, Time: p=0.008; 2W-ANOVA, Condition: p=0.88, Time: p=0.47, Interaction: p=0.89). (**B**) Percentage increase of sEPSC frequency from P21 to P42 and P42 to P85 for both CTL and SPD slices (P21-P42 T-Test, p=0.95; P42-P85 T-Test, p=0.87). (C-E) Amplitude (**C**) (CTL 1W-ANOVA, Time: p=0.038; SPD 1W-ANOVA, Time: p=0.054; 2W-ANOVA, Condition: p=0.57, Time: p=0.009, Interaction: p=0.84), Rise time (**D**) (CTL 1W-ANOVA, Time: p=0.083; SPD 1W-ANOVA, Time: p<0.001; 2W-ANOVA, Condition: p=0.24, Time: p<0.001, Interaction: p=0.048) and Decay time (**E**) (CTL 1W-ANOVA, Time: p=0.44; SPD 1W-ANOVA, Time: p=0.41; 2W-ANOVA, Condition: p=0.46, Time: p=0.30, Interaction: p=0.61) of sEPSC events. (**A-E**) Data from 8 (P21), 9 (P42 CTL), 11 (P42 SPD), 11 (P85 CTL), 9 (P85 SPD) cells. Statistical range: * p≤0.05; ** p<0.01; *** p<0.001

Intrinsic excitability of layer 5 pyramidal neurons was assessed in OFC slices from CTL and SPD rats. Similar to what was observed in the mPFC, the intrinsic excitability of OFC cells decreased from P21 to P42, but then recovered at P85 (Fig. 6A). AP threshold increased between P21 and P42 after which a small decrease was found at P85 (Fig. 6B), eventually coming back at P21 levels. The membrane potential (Fig. 6C) and input resistance (Fig. 6D) did not change over this developmental period. In SPD slices, the transient reduction in intrinsic excitability between P21 and P42 was absent (comparing Fig. 6A and Fig. 6E, P42 CTL-SPD 2W-ANOVA, condition: p<0.001), and AP firing rates showed a gradual increase over development (Fig. 6E). AP firing rates at P85 were comparable between CTL and SPD slices (comparing Fig. 6A and Fig. 6E, P85 CTL-SPD 2W-ANOVA, condition: p=0.56). The membrane potential (Fig. 6B), AP threshold (Fig. 6C) and input resistance (Fig. 6D) of the recorded neurons were unaffected after SPD. These experiments show that SPD has a small, but transient, effect on the intrinsic excitability of L5 cells in the OFC.

**Fig. 6.**
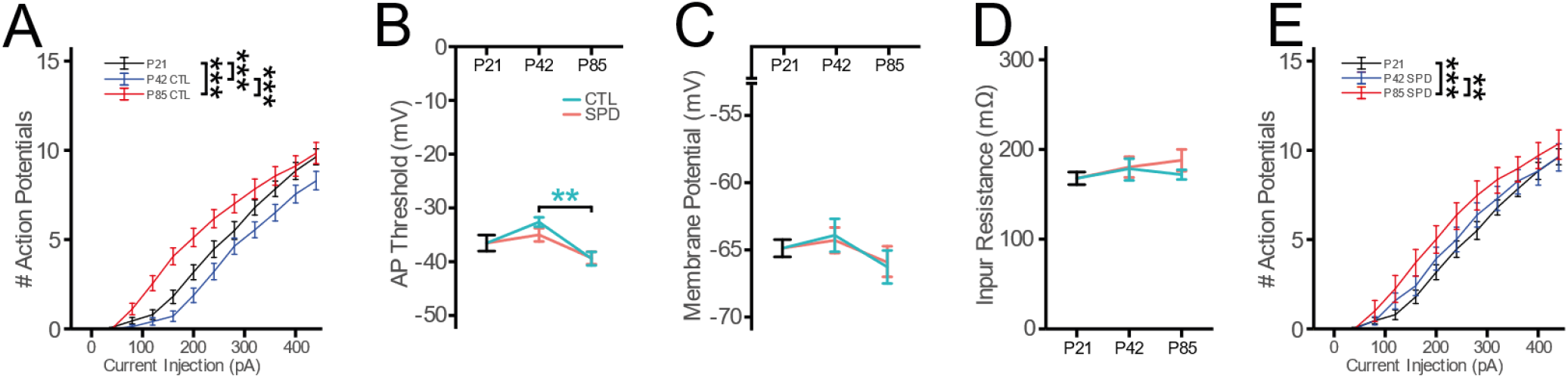
(**A**) Number of action potentials after current injections in Baseline (P21) and CTL (P42 & P85) OFC slices (2W-ANOVA, AP: p<0.001, Current: p<0.001, Interaction: p<0.001). (**B-D**) AP Threshold (**B**) (CTL 1W-ANOVA, Time: p=0.002; SPD 1W-ANOVA, Time: p=0.16; 2W-ANOVA, Condition: p=0.47. Time: p<0.001, Interaction: p=0.31), resting potential (**C**) (CTL 1W-ANOVA, Time: p=0.31; SPD 1W-ANOVA, Time: p=0.60; 2W-ANOVA, Condition: p=0.88, Time: p=0.11, Interaction: p=0.70) and input resistance (**D**) (CTL 1W-ANOVA, Time: p=0.74; SPD 1W-ANOVA, Time: p=0.46; 2W-ANOVA, Condition: p=0.51. Time: p=0.97, Interaction: p=0.59). (**E**) Number of action potentials after current injections in Baseline (P21) and SPD (P42 & P85) slices (2W-ANOVA, AP: p<0.001, Current: p<0.001, Interaction: p= 0.99). (**A-E**) Data from 24 (P21), 24 (P42 CTL), 27 (P42 SPD), 18 (P85 CTL), 13 (P85 SPD) cells. Statistical range: * p≤0.05; ** p<0.01; *** p<0.001

Together, these data highlight the differential development of synaptic connections onto L5 pyramidal cells in the mPFC and OFC. At P21, inhibitory and excitatory synaptic inputs onto L5 pyramidal cells were already present in the OFC, while these were largely absent, or at least silent, in the mPFC (Fig. 7A,B). SPD strongly affected the development of inhibitory inputs in both brain regions (Fig. 7D) while leaving excitatory synapses unaffected (Fig. 7D) and with only a transient effect on the firing properties of L5 cells. In both brain regions, inhibitory currents were reduced in adult slices from SPD rats, but this reduction occurred at different times in the OFC and mPFC. The strongest reduction of inhibitory inputs in the mPFC was observed immediately after SPD at P42, while the reduction of sIPSCs in the OFC occurred between P42 and P85.

**Fig. 7.**
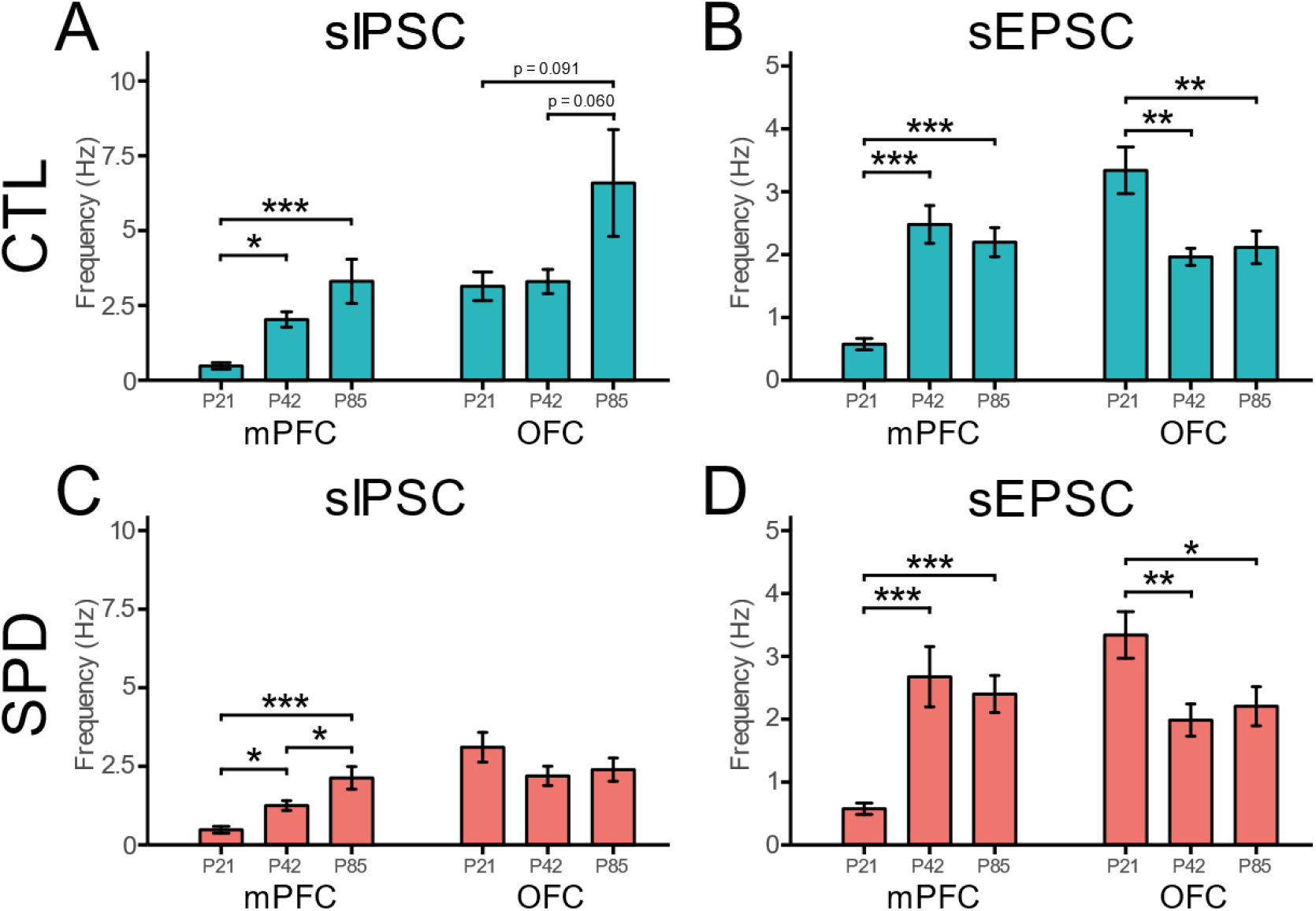
(**A,B**) Frequency of sIPSCs (**A**) (mPFC 1W-ANOVA, Time: p<0.001; OFC 1W-ANOVA, Time: p=0.049) and sEPSCs (**B**) (mPFC 1W-ANOVA, Time: p<0.001; OFC 1W-ANOVA, Time: p=0.002) in Baseline (P21) and CTL slices (P42 & P85) of both the mPFC and OFC. (**C,D**) Frequency of sIPSCs (**C**) (mPFC 1W-ANOVA, Time: p<0.001; OFC 1W-ANOVA, Time: p=0.265) and sEPSCs (**D**) (mPFC 1W-ANOVA, Time: p<0.001; OFC 1W-ANOVA, Time: p=0.007) in Baseline (P21) and SPD slices (P42 & P85) of both the mPFC and OFC. Statistical range: * p≤0.05; ** p<0.01; *** p<0.001

## Discussion

In this study we present the developmental timeline of inhibitory and excitatory synaptic inputs onto layer 5 pyramidal neurons in two subregions of the rat PFC, i.e. the mPFC and OFC. We found that these subregions develop with a differential time course and that SPD affects inhibitory but not excitatory inputs onto L5 pyramidal neurons in both regions, resulting in a specific reduction of inhibitory currents in adulthood. However, the reduction in IPSCs in the two PFC subregions arose via differential developmental trajectories. In the mPFC, development was mostly affected by SPD between P21 and P42, while IPSCs in the OFC were mainly affected after P42.

Social play enhances neural activity in the PFC and in corticostriatal and limbic structures which are connected to the PFC in the adult brain (Gordon et al., 2002, 2003; Hoover and Vertes, 2011; Van Kerkhof et al., 2014). Social play is almost absent before P21 (Baenninger, 1967; Panksepp, 1981), so activity in the PFC generated by social play is expected to be low at that age. During early adolescence (~P30 - P42), play is abundant and play-induced neural activity is likely one of the driving forces of PFC maturation. Both the OFC and mPFC have been implicated in social play behavior (Schneider and Koch, 2005; Van Kerkhof et al., 2013), but they exert a differential function in social interactions and cognitive flexibility. The mPFC in rats is important for shifting between cognitive strategies (Ragozzino et al., 1999; Birrell and Brown, 2000) and for coordination of movements during social interactions (Bell et al., 2010). In contrast, the OFC may be more involved in shifting between stimulus-reward associations (Ghods-Sharifi et al., 2008) and response modulation when interacting with different social play partners (Pellis et al., 2006). The current study shows that social play is one of the important driving forces of PFC maturation, and that SPD differentially affects the development of synaptic connections in the mPFC and the OFC.

To the best of our knowledge, our study is the first study that explicitly compares the developmental trajectory of inhibitory and excitatory synaptic inputs across these developmental timepoints in both the mPFC and OFC. So far, there have been a handful of studies that examined the development of postsynaptic inputs in the mPFC (Caballero et al., 2014b; Cass et al., 2014; Miyamae et al., 2017; Kroon et al., 2019; Kalemaki et al., 2020), but the development of the OFC circuitry has not been addressed. Our data show that the synaptic connections onto L5 cells in the mPFC and OFC develop via distinct trajectories (Fig. 7A,B). In the mPFC, the frequency of inhibitory inputs increases across juvenile and adolescent development with a strong increase in activity-dependent currents between P21 and P42. This coincides well with the described transition from an inhibitory system dominated mostly by regular spiking (calretinin (CR)-positive) to fast-spiking (parvalbumin (PV)-positive) interneurons (Caballero et al., 2014a; Caballero and Tseng, 2016) and the increasing excitatory drive onto PV interneurons (Caballero et al., 2014a). The modest additional increase in sIPSC frequency between P42 and P85 is also in agreement with a previous report (Cass et al., 2014). In contrast, the frequency of inhibitory inputs in the OFC was already high at P21 and remained stable until P42 (Fig. 7A). Comparison of mIPSC and sIPSC frequencies (Figs 1 and 4) indicates a strong activity-dependent contribution to the inhibitory drive, which occurs earlier in the OFC than in the mPFC. This suggests that the transition of the inhibitory system to a PV interneuron-dominated system occurred earlier in the OFC than in the mPFC. The development of excitatory inputs to L5 cells increased in the mPFC between P21 and P42 (Caballero et al., 2014a), but decreased in the OFC to reach comparable values at P42 (Fig. 7B). In both regions, sEPSC frequency remained stable after P42 until adulthood. Together, our data demonstrates a clear difference in the development of synaptic circuitry in these two main subregions of the rat PFC, which likely influences how they are affected by early life experience.

The mPFC and OFC are reciprocally connected, which was shown by extensive anatomical studies using antero- and retrograde tracers (Vertes, 2004; Hoover and Vertes, 2007, 2011). It is therefore likely that a synaptic change in one of the regions will affect the circuit development in the other. Consistent with our previous findings (Bijlsma et al., 2022), we observed that SPD affects the inhibitory, but not the excitatory, connections in both PFC regions. SPD resulted in reduced synaptic inhibition, reminiscent of the impaired development of inhibitory connections that has been described after sensory deprivation (Mowery et al., 2019; Reh et al., 2020). The reduction in IPSCs occurred before P42 in the mPFC, while inhibitory currents in the OFC were only affected after P42. This late effect in the OFC could either reflect a specific effect of the recovery from SPD in the OFC, or an indirect consequence of the reduced IPSCs in the mPFC and possibly other regions. It will be important to determine which inputs to the OFC are responsible for the difference in excitatory and inhibitory drive of L5 cells in the mPFC and OFC at P21, before the onset of play.

Together, our results demonstrate that excitatory and inhibitory synaptic inputs in the mPFC and OFC follow distinct developmental trajectories, and that lack of social play experience disturbs this development in a region-specific manner. This study highlights the differential vulnerability of PFC subregions to developmental insults, such as the lack of social play, which likely contributes to the multifaceted impact on cognitive performance in adulthood.

## Acknowledgements

This work was supported by the Netherlands Organisation for Scientific Research (NWO) (ALWOP.2015.105).

## Notes

### Competing Interest Statement

The authors have declared no competing interest.

